# Hippocampal Transcriptome Profiling Reveals Common Disease Pathways in Chronic Hypoperfusion and Ageing

**DOI:** 10.1101/2020.11.21.392449

**Authors:** Sang-Ha Baik, Sharmelee Selvaraji, David Y. Fann, Dong-Gyu Jo, Deron R. Herr, Mitchell K.P. Lai, Christopher Li-Hsian Chen, Grant R. Drummond, Kah-Leong Lim, Christopher G. Sobey, Thiruma V. Arumugam

## Abstract

Vascular dementia (VaD) is a progressive cognitive impairment of vascular etiology. VaD is characterized by cerebral hypoperfusion, increased blood-brain barrier permeability and white matter lesions. An increased burden of VaD is expected due to rapidly aging populations. The hippocampus is particularly susceptible to hypoperfusion, and the resulting memory impairment may play a crucial role in VaD. Here we have investigated the hippocampal gene expression profile of young and old mice subjected to chronic cerebral hypoperfusion by bilateral common carotid artery stenosis (BCAS). Our data in sham-operated young and aged mice show the normal age-associated decline in cerebral blood flow and differential gene expression. BCAS and ageing caused broadly similar effects, however, BCAS-induced changes in hippocampal gene expression differed between young and aged mice. Specifically, transcriptomic analysis indicated that in comparison to young sham mice, many pathways altered by BCAS in young mice resembled those present in sham aged mice. Immunoblot analyses confirmed these findings. Finally, relative to young sham mice the cell type-specific profile of genes in both young BCAS and old sham animals further revealed common cell-specific genes. Our data provide a genetic-based molecular framework for chronic hypoperfusion-induced hippocampal damage and reveal common cellular signaling pathways likely to be important in the pathophysiology of VaD.

## Introduction

Vascular dementia (VaD), like Alzheimer’s disease, is a common cause of dementia (1) where there is an increased risk with aging, rapid, step-wise disease progression and a high mortality rate (2). While the root cause of the problem remains unknown, the known risk factors driving VaD include diabetes, hypertension and metabolic syndrome (1). The current treatment options for VaD, are therefore, are purely symptomatic which heightens the urgency to better understand and treat this condition at its roots. Both clinical and experimental data indicate that cerebrovascular disease, as well as cardiac and systemic vascular diseases, may lead to cerebral hypoperfusion, and thus altering brain metabolism and leading to cognitive impairment (4, 5, 7, 8). Indeed, cerebral hypoperfusion-induced microscopic infarcts (microinfarcts) and white matter injuries are common features in VaD patients (8). However, the mechanisms by which cerebral hypoperfusion-induced microinfarcts promote VaD remain unclear (9-12).

In order to develop a mechanistic understanding of VaD, this study employs a chronic cerebral hypoperfusion mouse model of VaD:bilateral common carotid artery stenosis (BCAS) model. It is a widely used animal model of VaD that induces subcortical ischemia (1,13). BCAS employs external microcoils around both common carotid arteries in rodents to reduce cerebral blood flow (CBF) and hence induce damage to the deep white matter. BCAS closely mimics several features of VaD in patients, including decreased metabolism, elevated inflammation, decreased blood brain barrier (BBB) integrity, white matter lesions, neuronal damage and working memory deficits (13, 14-17). In particular, CBF to the hippocampus, a limbic structure pivotal in learning and memory and vulnerable to chronic stress (19), is reduced following BCAS (18), which is consistent with hippocampal atrophy in patients with subcortical VaD (14). However, the transcriptional processes underlying this hippocampal susceptibility in VaD remain unknown. The present study has thus investigated the hippocampal gene expression profile of young and old mice subjected to sham surgery or BCAS for up to 30 days.

## Results

### BCAS reduces cerebral blood flow and alters gene expression in the brain of young and old mice

Young (3 months old) and old (22 months old) male C57BL/6 mice were randomly assigned to receive either Sham or BCAS surgery. CBF was measured at baseline, immediately after Sham/BCAS surgery, and prior to euthanasia after 7 or 30 days (**Fig. 1A and B**). CBF in all groups was expressed relative to mean CBF perfusion in young sham-operated mice at baseline. An age-associated decline in CBF was observed in old versus young sham-operated mice (**Fig. 1B and C**). BCAS reduced CBF by ~30% in both young and old mice (**Fig. 1B and C**). Interestingly, there was no difference in blood flow observed between (a) old Sham and young BCAS mice and (b) old and young BCAS mice at both 7 and 30 day time points (**Fig. 1B and C**). This highlights the overlap between the age-associated and disease-associated decline in CBF.

**Figure 1:**
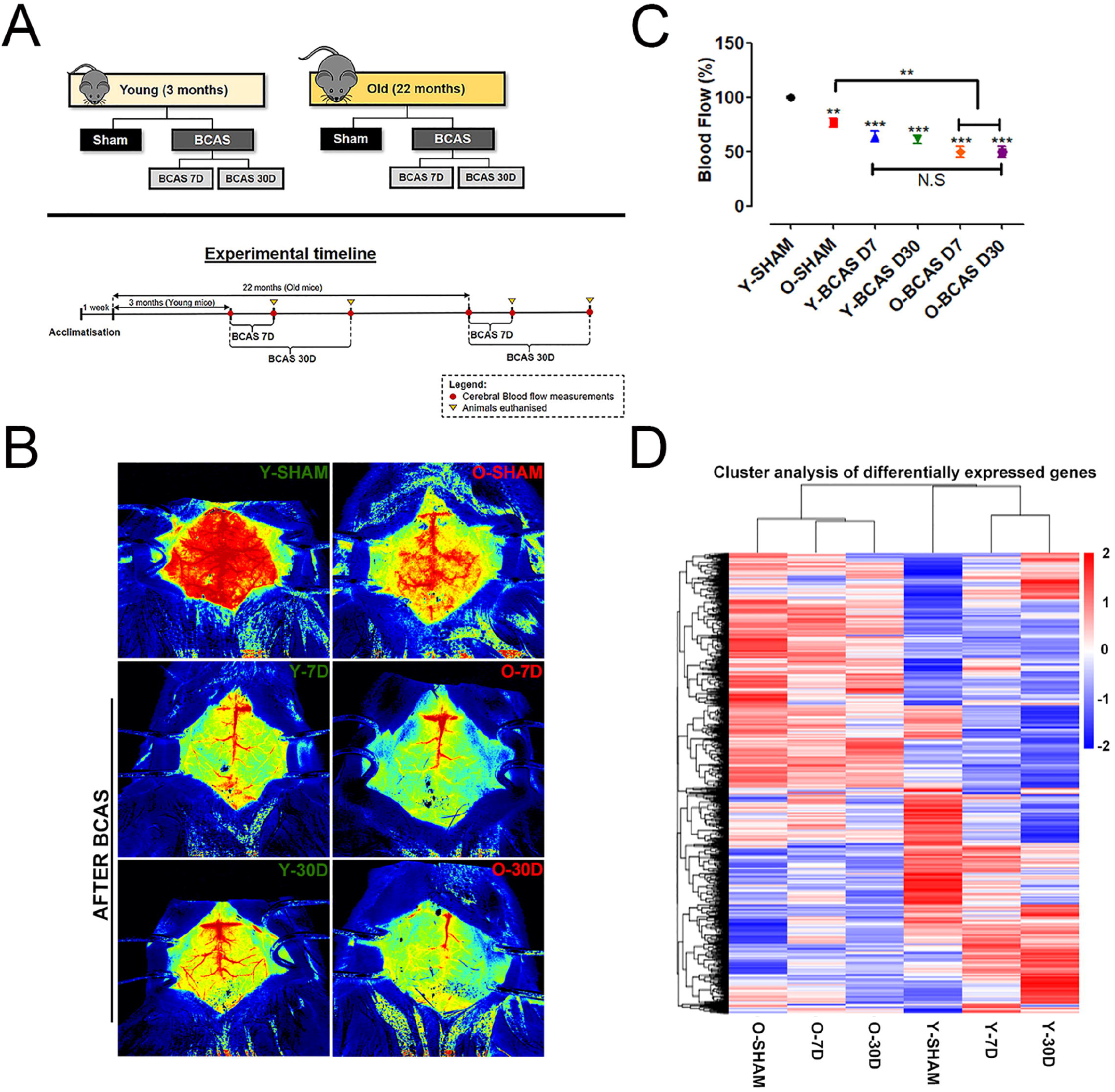
Effect of bilateral common carotid artery stenosis (BCAS) on cerebral blood flow and gene expression in young and old mice. (**A**) Experimental model and timelines of blood flow measurements and tissue collection. Laser speckle contrasting imaging was used to monitor the cerebral blood perfusion in real time for all experimental groups before and after insertion of microcoils through BCAS surgery. (**B & C**) Representative contrast images and quantification of basal cerebral blood flow before surgery, and effective blood flow reduction after 7 and 30-days of BCAS. The rate of blood flow was calculated as an absolute value of cerebral perfusion in perfusion units (PU) using the PeriMed Software. Data are represented as mean ± S.E.M. n = 5 mice in each experimental group. **P<0.01; ***P<0.001. (**D**) Heatmap of differentially expressed genes in Old Sham (OSham), Old 7 Day (O7D) BCAS, Old 30 Day (O30D) BCAS, Young Sham (YSham), Young 7 Day(Y7D) BCAS and Young 30 Day(Y30D) BCAS, groups with upregulated genes in red and downregulated genes in blue. The colour scale represents the log 10 (FPKM□+□1) value.

Unsupervised hierarchical clustering of differentially expressed genes (DEGs) in the hippocampus revealed significant differences between young and old sham-operated mice (**Fig. 1D**). Heatmap patterns also varied in a time-dependent manner over 30 days post-BCAS surgery, as indicated by clustering (see below). It is notable that the DEG heatmap patterns of OSham are similar to that of O7D and O30D BCAS mice.

### Ageing

There were 1688 significant DEGs (898 upregulated and 790 downregulated) between young and old sham-operated mice (**Fig. 2A**). We annotated these age-associated DEGs by performing differential enrichment analysis using cluster profiling. Transcriptome analysis using gene ontology (GO) enrichment showed that the top 20 significantly upregulated GO terms in the old compared to young mice belonged to a myriad of development and cell migration processes, including angiogenesis, morphogenesis, cell adhesion, tissue migration and response to external stimulus (**Fig. 2B; Supplementary Table 1**). Upregulation of GO terms such as tissue migration and antigen processing and presentation of peptide antigen also suggest upregulated immunological processes. The top 20 downregulated enriched GO terms for the old versus young mice belonged to physiological functions such as cell redox homeostasis, mitochondrial transmembrane transport, and key metabolic processes such as sulfur compound metabolic process, nucleoside triphosphate metabolic processes, ATP metabolic process, purine ribonucleoside metabolic processes, cellular respiration and protein folding (**Fig. 2B**).

**Figure 2:**
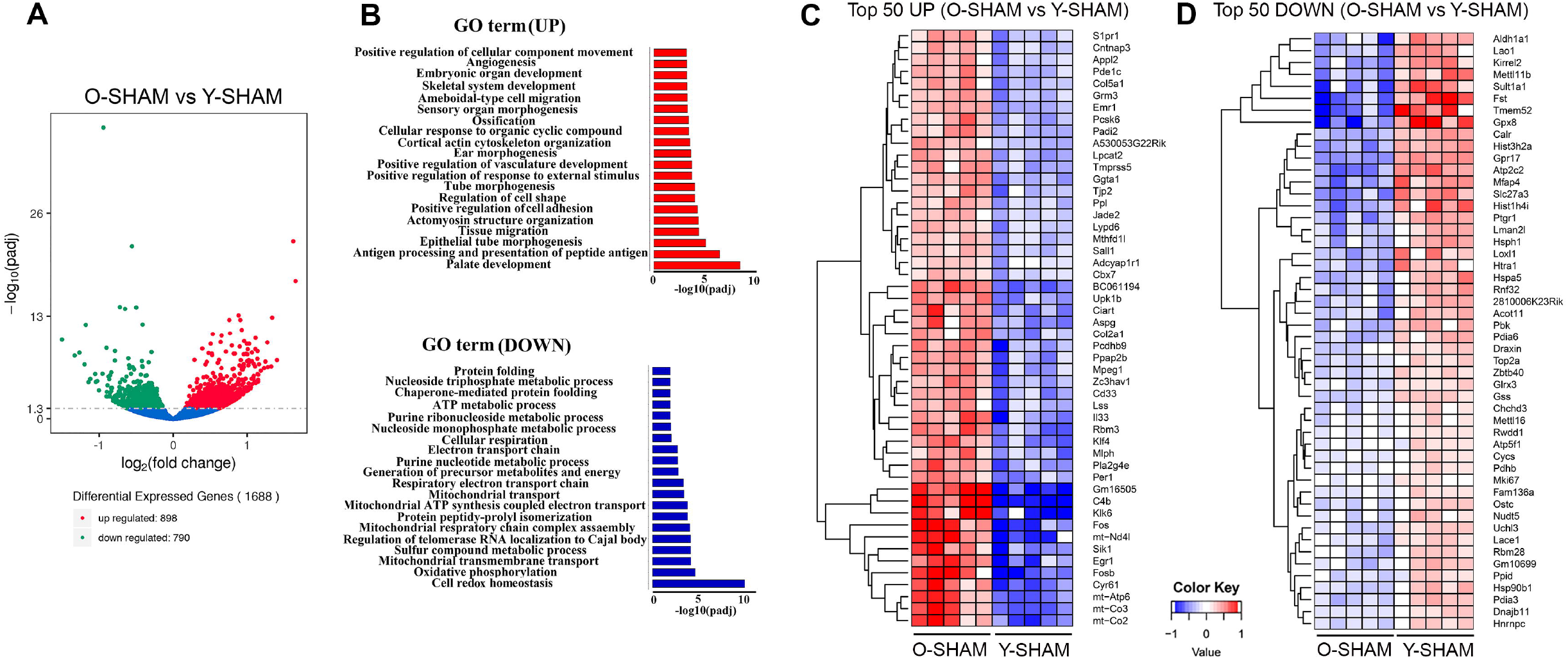
Analysis of differentially expressed genes in young and old hippocampus and their enriched GO terms. (**A**) Volcano plot of differentially expressed genes analyzed in old Sham group against young Sham group. The threshold of differential expression is q-value < 0.05. The horizontal axis is the log2 fold change of genes. The vertical axis is statistical significance scaled as −log 10 q-value. Each dot represents an individual gene (blue: no significant difference; red: upregulated expression of gene; green: down-regulated expression of gene). (**B**) The selection of significantly enriched top upregulated and down-regulated GO terms are presented for old Sham against young Sham animals (q < 0.05). The horizontal axis represents the −log10 scale for q-value of each GO term. (**C**) Top 50 up-regulated genes in old Sham against young Sham animals. (**D**) Top 50 down-regulated genes in old Sham against young Sham animals. Up-regulated genes in red and downregulated genes in blue. The color scale represents the log10 (average FPKM+0.5) value.

The top 50 upregulated genes in the old compared to young mice include mitochondrially encoded ATP synthase membrane subunit 6 (*Atp6*), mitochondrially encoded cytochrome C oxidase III (*mt-Co3*), mitochondrially encoded cytochrome C oxidase II (*mt-Co2*), mitochondrially encoded NADH 4L dehydrogenase (*mt-nd4l*), genes involved in inflammation such as cysteine-rich angiogenic inducer 61 (*Cyr61*), early growth response 1 (*Egr1*), kallikrein related peptidase 6 (*Klk6*), salt inducible kinase 1 (*Sik1*), complement C4B (*C4b*), Kruppel Like Factor 4 (*Klf4*), apoptosis-related genes such as proto-oncogene, AP-1 transcription factor subunit (*Fos*) (20) and genes that regulate adult neurogenesis such as PH domain and leucine zipper 2 (*APPL2*) (21) (**Fig. 2C; Supplementary Table 2**). The top 50 downregulated genes include, aldehyde dehydrogenase 1 family member A1 (*Aldh1a1*), activin-binding follistatin (*Fst*) (22), anti-oxidative glutathione peroxidase 8 (*GPx8*) (23) and prostaglandin reductase 1 (*Ptgr1*) (24), sulfotransferase family 1A member 1 (*Sult1a1*) and ATPase secretory pathway Ca^2+^ transporting 2 (*Atp2c2*) that modulate phonological short-term memory in language impairment (25) and a gene encoding a member of a family of adenosine triphosphate (ATP)-metabolizing molecular chaperones with roles in stabilizing and folding other proteins, heat shock protein 90 beta family member 1 (*Hsp90b1*) (**Fig. 2D; Supplementary Table 2**).

### Cerebral Hypoperfusion

We assessed the effect of BCAS after 7 and 30 days on hippocampal gene expression in young and old mice. In young mice, there were global changes of gene expression patterns at both time points and significant DEGs that deviated from that of young sham-operated mice (**Fig. 3A**). However, in old mice, cerebral hypoperfusion did not lead to profound changes in gene expression after either 7 or 30 days of BCAS compared to sham-operated old mice (**Supplementary Fig. 1**). We thus focused our analyses on transcriptomic changes after BCAS in young mice. After 7 days, young mice had 279 significantly upregulated and 299 significantly downregulated genes compared to sham mice (**Fig. 3B**), with more DEGs at 30 days than 7 days of BCAS. At 30 days, 598 genes were significantly upregulated and 552 genes were significantly downregulated in Y30D compared to sham mice (**Fig. 3B**).

**Figure 3:**
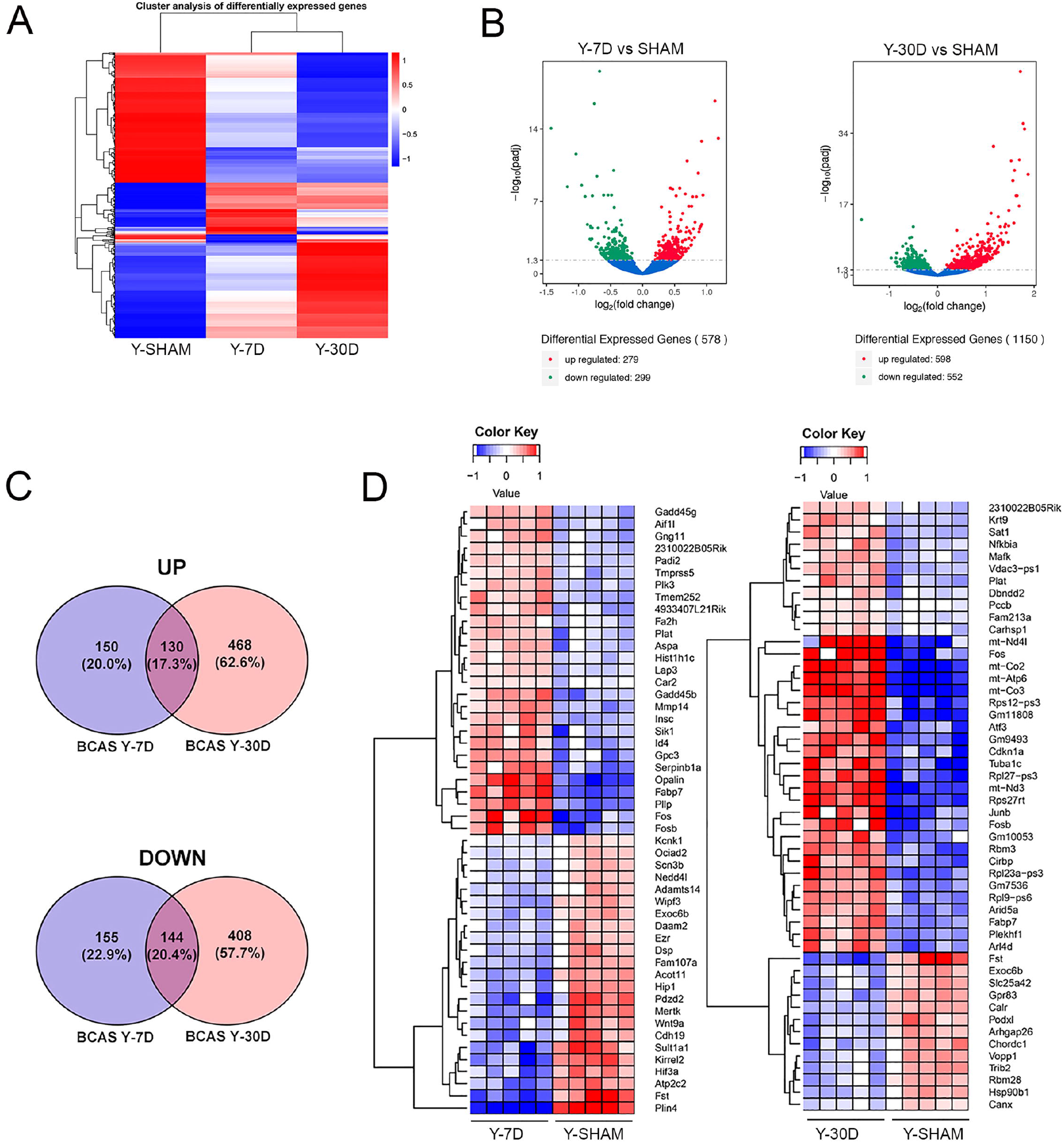
Differentially expressed genes in young hippocampus following chronic hypoperfusion. (**A**) Hierarchical clustering analysis of differentially expressed mRNA transcripts following 7 and 30 days BCAS compared to Sham animals. Upregulated genes in red and downregulated genes in blue. The color scale represents the log10 (average FPKM+1) value. (**B**) Volcano plots of differentially expressed genes analyzed in young 7 day and young 30 day BCAS animals compared to Sham animals. The threshold of differential expression is q value < 0.05. The horizontal axis is the log2 fold change of transcripts. The vertical axis is statistical significance scaled as −log 10 q-value. Each dot represents an individual transcript (blue: no significant difference; red: upregulated expression of gene; green: downregulated expression of gene). (**C**) Top 50 upregulated genes in young 7 day and young 30 day BCAS animals compared to Sham animals. (**D**) Venn diagram reflects the distribution of differentially expressed genes from 7 day and 30 day BCAS and Sham comparisons.

A full list of DEGs in the 7 and 30 days of BCAS groups is provided in **Supplementary Table 3**. The results of GO term analysis in young mice subjected to 7 days of BCAS revealed upregulation of genes involved in biological processes such as cell substrate junction, focal adhesion, myelin sheath, extracellular matrix, response to toxic substance, while biological processes such as ion channel complex, ion channel activity, transmembrane transporter complex, calcium channel activity, plasma membrane protein complex were significantly downregulated (**Supplementary Fig. 2**). In addition to GO enrichment analysis, we performed Kyoto Encyclopedia of Genes and Genomes (KEGG) pathway enrichment analysis of the RNA sequencing data. Those results were generally similar to those of the GO term analysis, indicating pathways involved in focal adhesion, chemokine signaling pathway and long-term depression. KEGG analysis also revealed downregulation of gene sets related to calcium signaling, regulation of actin cytoskeleton, axon guidance, and memory (**Supplementary Fig. 3 and Supplementary Table 4**).

The results of GO analysis revealed upregulation of genes involved in transcription factor activity, cofactor binding, mitochondrial inner membrane, organelle inner membrane, regulation of cellular amide metabolic process, cellular response to growth factor stimulus and reactive oxygen species metabolic process after 30 days of BCAS in young mice. GO terms such as regulation of cell morphogenesis, protein serine/threonine kinase activity, post-synaptic function and density, neurospine, excitatory synapse, dendritic spine, axon extension, ion channel complex, phospholipid binding, transporter complex and transmembrane transporter complex were downregulated in these mice (**Supplementary Fig. 4 and Supplementary Table 5**).

### Temporal changes in DEGs during chronic cerebral hypoperfusion

The spread of unique DEGs after 7 or 30 days of BCAS in young mice were summarized in a Venn diagram (**Fig. 3C**). There were some DEGs in common at 7 and 30 days, with 130 genes upregulated and 144 genes downregulated (**Supplementary Table 6**); however, there were approximately 3 times more DEGs in either direction at 30 versus 7 days (**Fig. 3C**), consistent with a general temporal progression of hypoperfusion-induced injury at the transcriptome level (**Fig. 3C**).

We narrowed our investigation to the top 50 DEGs at 7 and 30 days post-BCAS (**Fig. 3D**). At 7 days, downregulated DEGs included a lipid marker of neurodegeneration transcript perilipin 4 (*Plin4*) (26), follistatin (*Fst*) (22), protein coding genes sulfotransferase family 1A member 1 (*Sult1a1*) and ATPase secretory pathway Ca^2+^ transporting 2 (*Atp2c2*), hypoxia inducible factor 3 subunit alpha (*Hif3a*), which acts as a transcriptional regulator in adaptive response to low oxygen tension (27) and myelin-forming cell-specific cadherin 19 (*Cdh19*) (28) among many other genes listed in **Figure 3D**. Upregulated DEGs included death promoting FosB proto-oncogene, AP-1 transcription factor subunit (*Fosb*) and Fos proto-oncogene, AP-1 transcription factor subunit (*Fos*) (29, 30), plasma membrane proteolipid (*Pllp*), and oligodendrocytic myelin paranodal and inner loop protein (*Opalin*) (**Fig. 3D**).

At 30 days post-BCAS, downregulated DEGs included follistatin (*Fst*) (22), exocyst complex component 6B (*Exoc6b*), mitochondrial coenzyme A transporter (*Slc25a42*), stress response gene G Protein-Coupled Receptor 83 (*Gpr83*) (31), Ca^2+-^binding chaperone protein Calreticulin (*Calr*) and neuroprotective heat shock protein 90kDa beta (*Grp94*) and member 1 (*Hsp90b1*) (32). Upregulated DEGs included genes that were also upregulated in both old Sham mice and young mice at 7 days post-BCAS. These genes included: AP-1 transcription factor subunit (*Fos*), mitochondrially encoded cytochrome C oxidase III (*mt-Co3*), mitochondrially encoded cytochrome C oxidase II (*mt-Co2*), ATP synthase membrane subunit 6 (*Atp6*), mitochondrially encoded NADH:Ubiquinone oxidoreductase core subunit 3 (mt-nd3), cyclin dependent kinase inhibitor 1A (Cdkn1a), FosB proto-oncogene and AP-1 transcription factor subunit (*Fosb*) (29, 30) (**Fig. 3D**).

### Cerebral hypoperfusion and ageing exhibit common changes in the hippocampal transcriptome

We assessed hippocampal transcriptome data from old sham-operated mice and young mice subjected to BCAS for 30 days in comparison with young sham-operated mice (**Fig. 4**). The heatmap shows global changes of gene expression in these groups (**Fig. 4A**). We found that over 4000 genes were differentially expressed between old sham-operated mice and young mice subjected to BCAS for 30 days with over 2300 genes downregulated (**Fig. 4A**). The GO term analysis revealed that compared to BCAS-operated mice, old sham mice had downregulated biological processes mainly related to mitochondrial integrity (e.g. mitochondrial ribosome, mitochondrial inner membrane, mitochondrial protein complex, respiratory chain, mitochondrial matrix and NADH dehydrogenase complex) (**Fig. 4C**). It is noteworthy that Venn diagram analyses indicated that old sham-operated mice and young mice subjected to BCAS for 30 days had in common 117 genes that were upregulated and 110 genes that were downregulated (**Fig. 4D-4E**). The common DEGs include those involved in the mitochondrial respiratory chain complex (**Fig. 4F**) (e.g. mitochondrially encoded NADH dehydrogenase 3 and 4, *Mt-Nd3; Mt-Nd4;* mitochondrially encoded cytochrome c oxidase II and III, *Mt-Co2; Mt-Co3;* and mitochondrially Encoded ATP Synthase Membrane Subunit 6, *Mt-Atp6*. The 110 genes downregulated in both groups included mitochondrial genes (e.g. NADH:Ubiquinone Oxidoreductase Subunit B2 and NADH:Ubiquinone Oxidoreductase Complex Assembly Factor 4, *Ndufaf4*) (**Fig. 4F**) and neuroprotective heat shock proteins (e.g. heat shock protein family H, *Hsp110*; member 1, *Hsph1*; heat shock protein family A, *Hsp70*; member 5, *Hspa5*; heat shock protein 90 alpha family class A member 1, *Hsp90aa1*; and heat shock protein 90 beta family member 1, *Hsp90b1*) (**Fig. 4G**).

**Figure 4:**
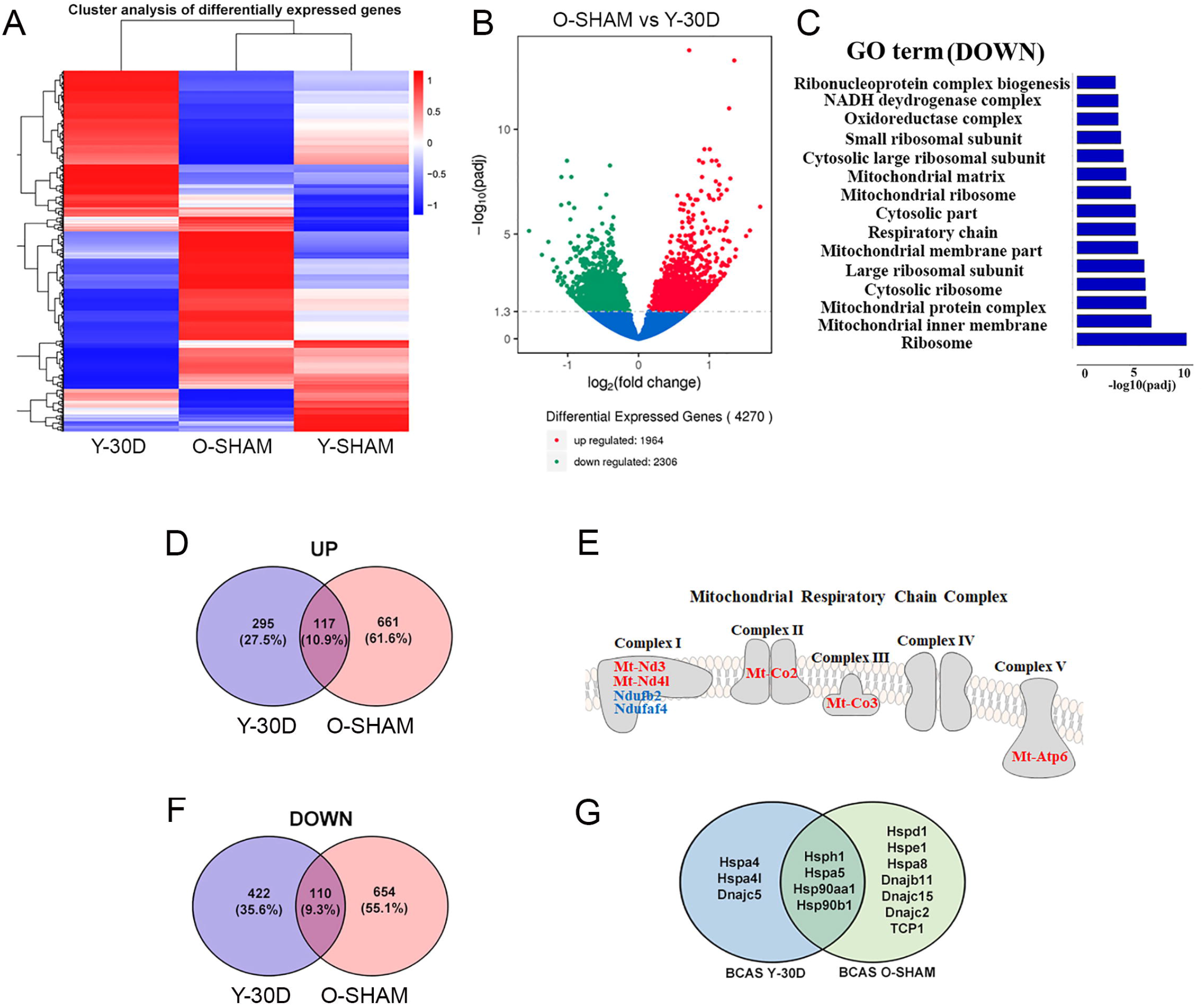
Comparative analyses of differentially expressed genes between young BCAS hippocampus and old Sham. (**A**) Hierarchical clustering analysis of differentially expressed genes following young 30 days BCAS and old Sham. Upregulated genes in red and down-regulated genes in blue. The color scale represents the log10 (average FPKM+1) value. (**B**) Differentially expressed mRNA transcripts analyzed between young 30 day BCAS and old Sham animals. The threshold of differential expression is q value < 0.05. The horizontal axis is the log2 fold change of transcripts. The vertical axis is statistical significance scaled as −log 10 q-value. Each dot represents an individual transcript (blue: no significant difference; red: upregulated transcript; green: down-regulated transcript). (**C**) The selection of significantly enriched down-regulated GO terms in both young 30 day BCAS and old Sham animals (q < 0.05). The horizontal axis represents the −log10 scale for q-value of each GO term. (**D**) Venn diagram reflects the distribution of differentially upregulated and downregulated genes from young 30-day BCAS and old Sham comparisons. (**E**) Venn diagram reflects the distribution of differentially down-regulated mRNA transcripts from young 30-day BCAS and old Sham comparisons. (**F**) Common upregulated (red) and downregulated (blue) mitochondrial genes in both young 30-day BCAS and old Sham hippocampus. (**G**) Common and unique down-regulated protein chaperons in young 30 day BCAS and old Sham groups.

Our observation that ageing and chronic cerebral hypoperfusion may both lead to hippocampal mitochondrial dysfunction and protein dysregulation led us to next validate our data by examining protein expression levels of selected genes (**Fig. 5**). Immunoblot analysis confirmed that compared to young sham mice, mitochondrial proteins such as Mt-Atp6, Mt-Co2, Mt-Co3 were upregulated in old sham mice, while expression of Mt-Atp6 and Mt-Co2 tended to be higher in young mice subjected to 30 days of BCAS (**Fig. 5A**). Neuroprotective protein chaperones such as GRP78 and Calnexin were downregulated following BCAS, while GRP78 was downregulated in old sham mice (**Fig. 5B**).

**Figure 5:**
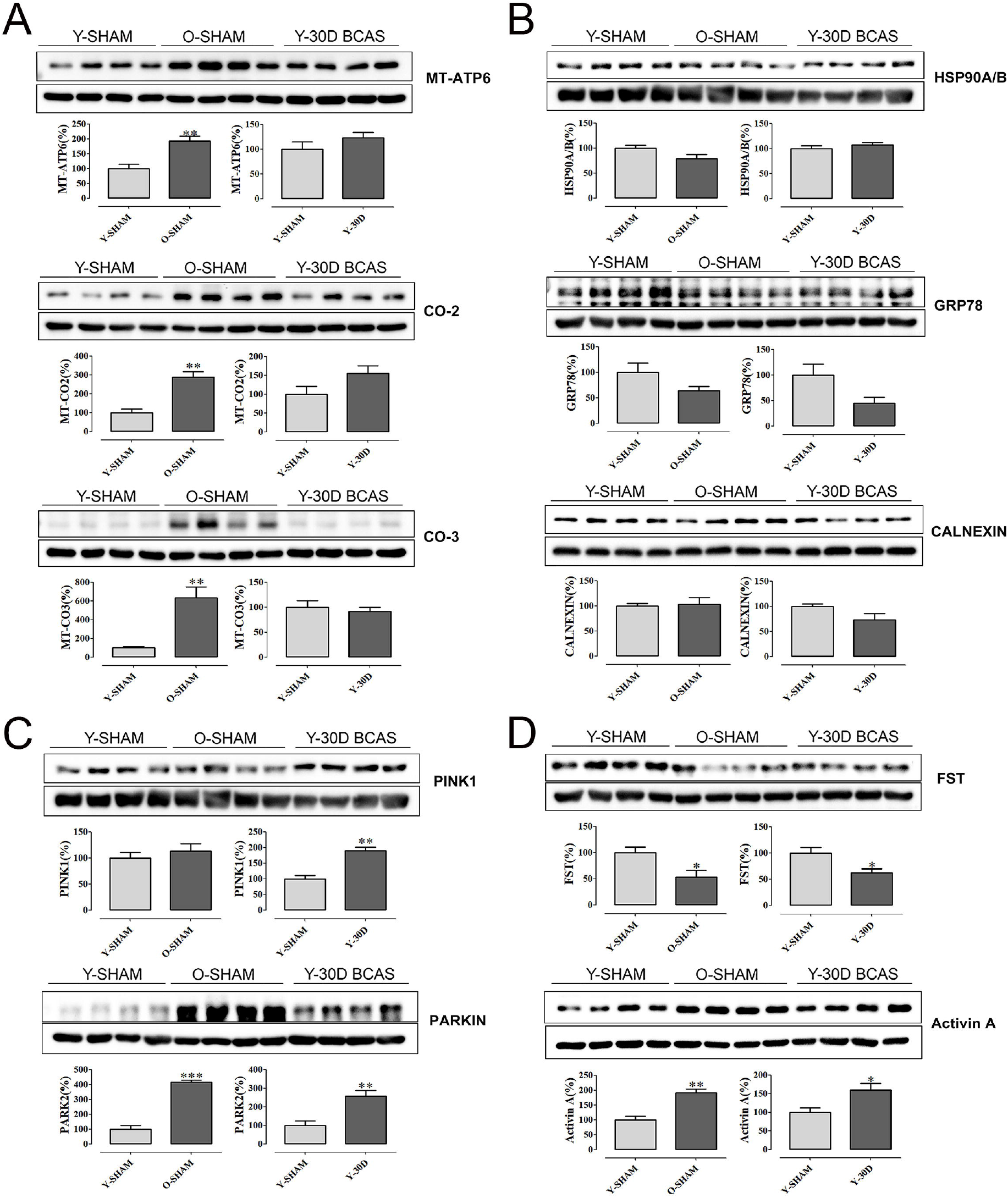
Immunoblot validation of mRNA transcripts. (**A**) Selected mitochondrial proteins Mt-Atp6, Mt-Co2, Mt-Co-3 and (**B**) protein chaperons Hsp90A/B, Grp78, Calnexin and (**C**) mitochondrial quality control proteins Park2 & Pink1 and (**D**) neuroprotective Activin A and its antagonist Fst. Data are represented as mean ± S.E.M. n=5-7 mice in each experimental group. *P<0.05 compared with young Sham; **P<0.01 compared with young Sham; ***P<0.001 compared with young Sham.

To further examine the common pathways potentially altered in both ageing and VaD pathology, we investigated the expression levels of PTEN-induced kinase 1 (Pink1) and E3 ubiquitin protein ligase PARKIN, two proteins which govern mitochondrial quality control (33). Compared to young sham mice, hippocampal expression of both proteins was significantly higher after cerebral hypoperfusion for 30 days, and PARKIN was also higher in old sham mice (**Fig. 5C**). We also validated two other proteins, Activin A and its antagonist follistatin (FST) (22), genes for which were differentially expressed in both old sham mice and young mice following 30 days of BCAS. Our immunoblot data followed the patterns of gene expression, whereby FST was lower in these groups compared to young sham mice (**Fig. 5D**). By contrast, Activin A expression was higher in these two treatment groups compared to young sham mice (**Fig. 5D**).

### Cell-type specific analyses

We next sorted the DEGs according to cell-type specificity based on previously published databases (34-37). Relative to young sham mice, the cell-type specific profiles of significant DEGs are represented in six separate Venn diagrams according to the following: astrocytes, endothelial cells, microglia, neurons, oligodendrocytes and oligodendrocyte progenitor cells (OPCs) (**Fig. 6A**). Protein-protein interaction (PPI) analyses revealed that most of the proteins associated with DEGs were independent of each other (**Fig. 6B-E**).

**Figure 6:**
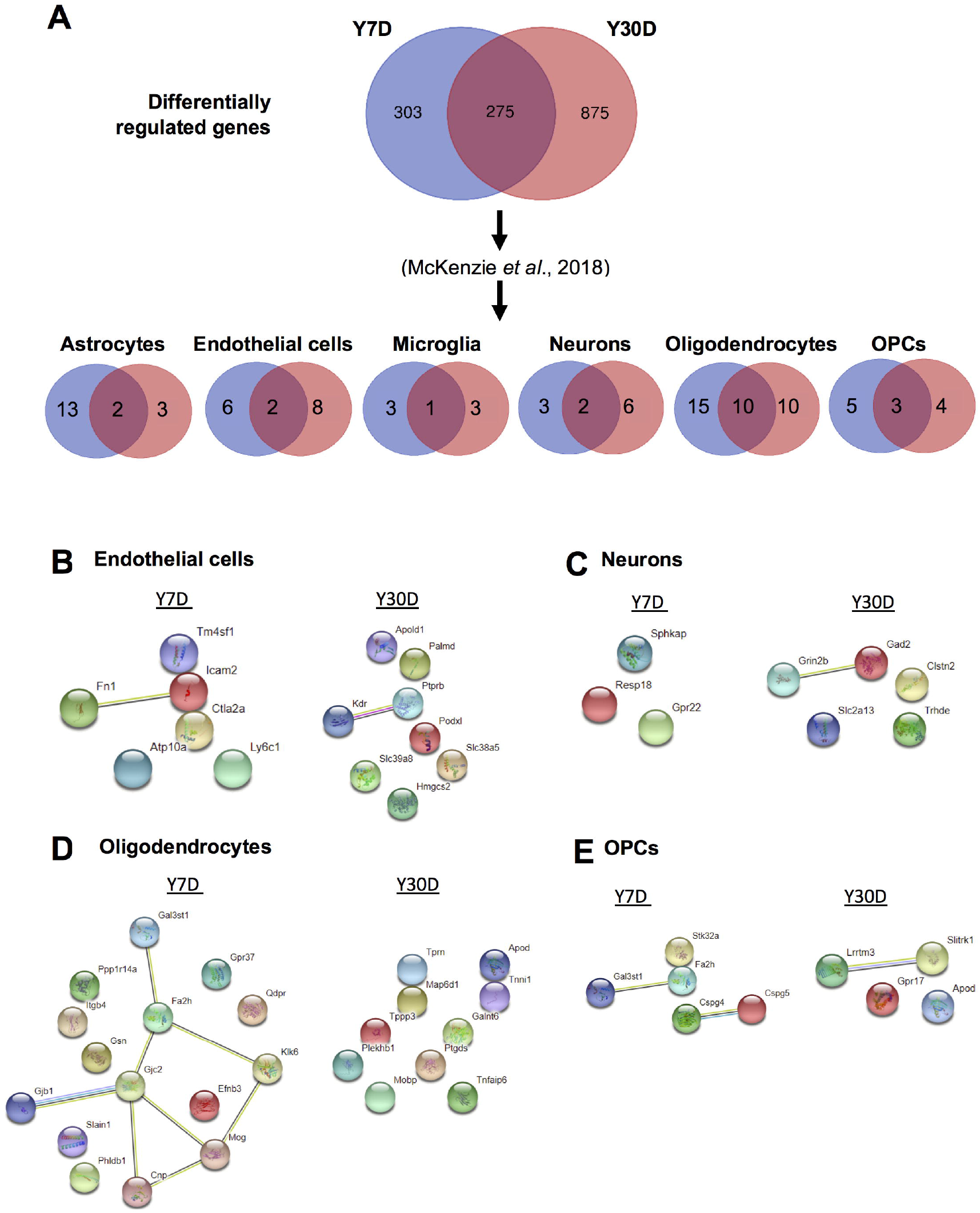
Analyses of differentially expressed mRNA transcripts in different brain cells following BCAS in young animals. **(A)** Venn diagram of statistically significant (adjusted p-value<0.05) differentially regulated genes in the Y7D/YSham vs Y30D/YSham comparison groups. The data was sorted according to cell type specificity based on a previously published database (Ref. 34) (Refer to Materials & Methods section for more details). The cell type specific profile of the genes are illustrated in six venn diagrams for each cell type including astrocytes, endothelial cells, microglia, neurons, oligodendrocytes and oligodendrocyte precursor cells (OPCs). The potential protein-protein interactions of cell type specific genes unique for the respective experimental groups of Y7D and Y30D respectively for endothelial cells **(B)**, neurons **(C)**, oligodendrocytes **(D)** and OPCs **(E)**. The lines joining the different gene nodes indicate protein-protein associations such as known or predicted interactions and others such as text mining, co-expression or protein homology.

A similar analysis to identify age-associated cell-type specific genes that may promote disease pathology was then performed (**Fig. 7**). DEGs were sorted according to cell-type specificity and visualized in Venn diagrams (**Fig.7A**). PPI analyses identified prominent interactions between old mice and those subjected to BCAS for 30 days in each cell type (**Fig. 7B-G**). While most proteins associated with DEGs were independent of each other in most cell types, the proteins associated with DEGs in the microglia of old sham mice had extensive interactions associated with proteins that promote neuroinflammation (**Fig. 7D**).

**Figure 7:**
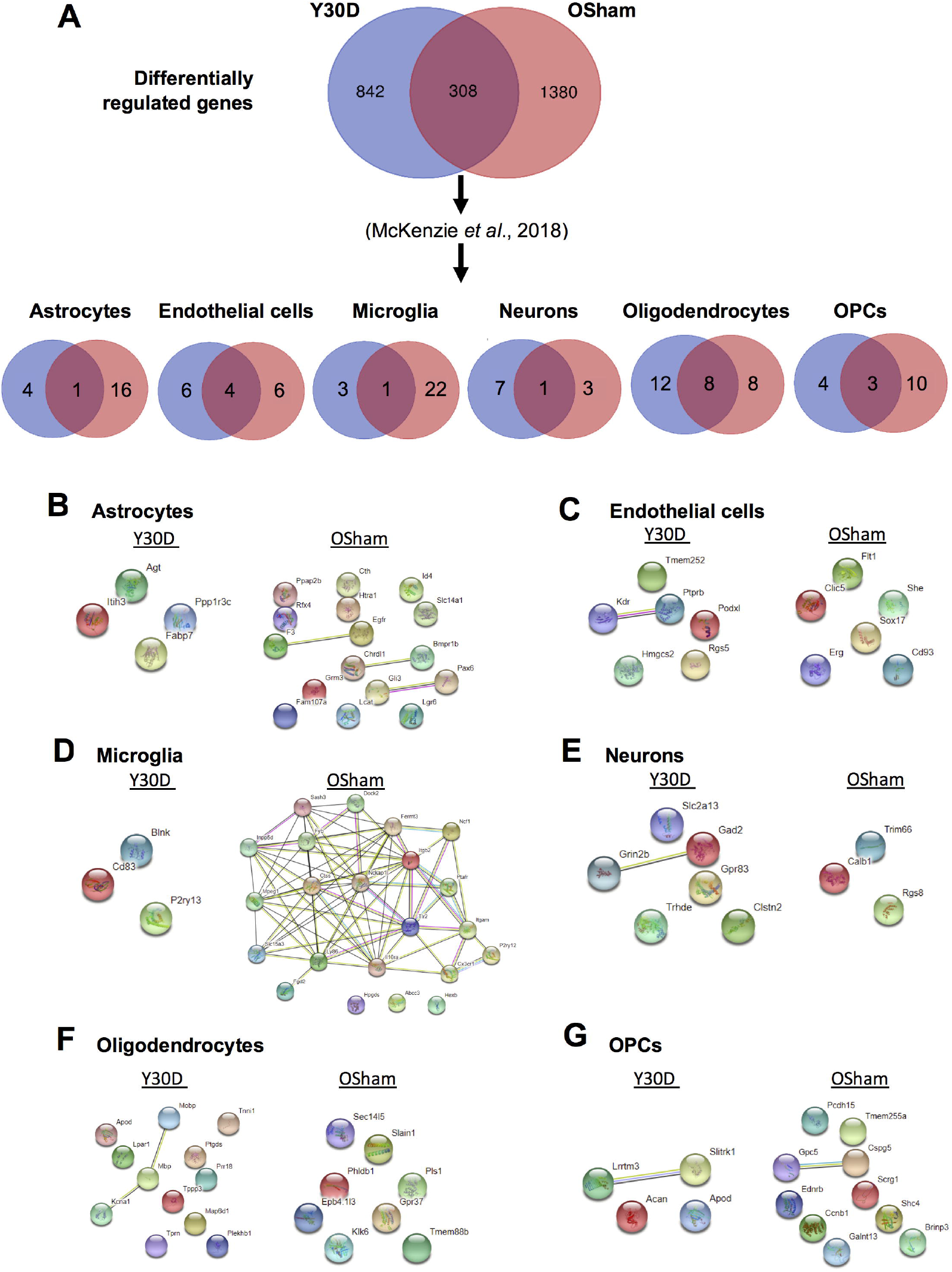
Analysis of differentially expressed mRNA transcripts in different brain cells in young 30 day BCAS animals compared to old Sham animals. **(A)** Venn diagram of statistically significant (adjusted p-value<0.05) differentially regulated genes in the Y30D/YSham vs OSham/YSham comparison groups. The data was sorted according to cell type specificity based on a previously published database (Ref. 34) (Refer to Materials & Methods section for more details). The cell type specific profile of the genes are illustrated in six venn diagrams for each cell type including astrocytes, endothelial cells, microglia, neurons, oligodendrocytes and oligodendrocyte precursor cells (OPCs). The potential protein-protein interactions of the cell type specific genes unique for the respective experimental groups of Y30D and OSham respectively for astrocytes **(B)**, endothelial cells **(C)**, microglia **(D)**, neurons **(E)**, oligodendrocytes **(F)** and OPCs **(G)**. The lines joining the different gene nodes indicate protein-protein associations such as known or predicted interactions and others such as text mining, co-expression or protein homology.

To further investigate the overlap in DEGs between old mice and those subjected to BCAS for 30 days, the associated cell-type specific data were analyzed (**Fig.8**). All gene expression values were normalized using the average of the respective control in young sham mice. The overlapping cell-type specific common genes as identified in the Venn diagrams (**Fig. 7A**) were Acyl-CoA Thioesterase 11 (*Acot11*) in astrocytes (**Fig. 8A**), solute carrier family 38 member 5 (Slc38a5), solute Carrier Family 39 Member 8 (*Slc39a8*), protein coding gene Palmdelphin (*Palmd*) and apolipoprotein L Domain Containing 1 (*Apold1*) in endothelial cells (**Fig. 8B**), lysophosphatidylcholine Acyltransferase 2 (*Lpcat2*) in microglia (**Fig. 8C**), calcium voltage-gated channel auxiliary subunit alpha2delta 3 (Cacna2d3) in neurons (**Fig. 8D**), arylsulfatase G (*Arsg*), aspartoacylase (*Aspa*), neurofascin (*Nfasc*), plasmolipin (*Pllp*), polypeptide N-acetylgalactosaminyltransferase 6 (*Galnt6*), INSC spindle orientation adaptor protein (*Insc*), oligodendrocytic myelin paranodal & inner loop protein (*Opalin*) and TNF alpha induced protein 6 (*Tnfaip6*) in oligodendrocytes (**Fig. 8E**), S100 calcium binding protein B (*S100b*) and G protein-coupled receptor 17 (*Gpr17*) in OPCs (**Fig. 8F**). While there was a general differential regulation of overlapping DEGs between old mice and those subjected to BCAS for 30 days across the different cell-types, it was notable that all but one of the DEGs classified under oligodendrocytes had relatively lower gene expression in old mice (**Fig.8E**). This cell-type specific analysis requires further validation to confirm the trends, but it serves as a starting point to identify overlaps between age- and disease-associated DEGs.

**Figure 8:**
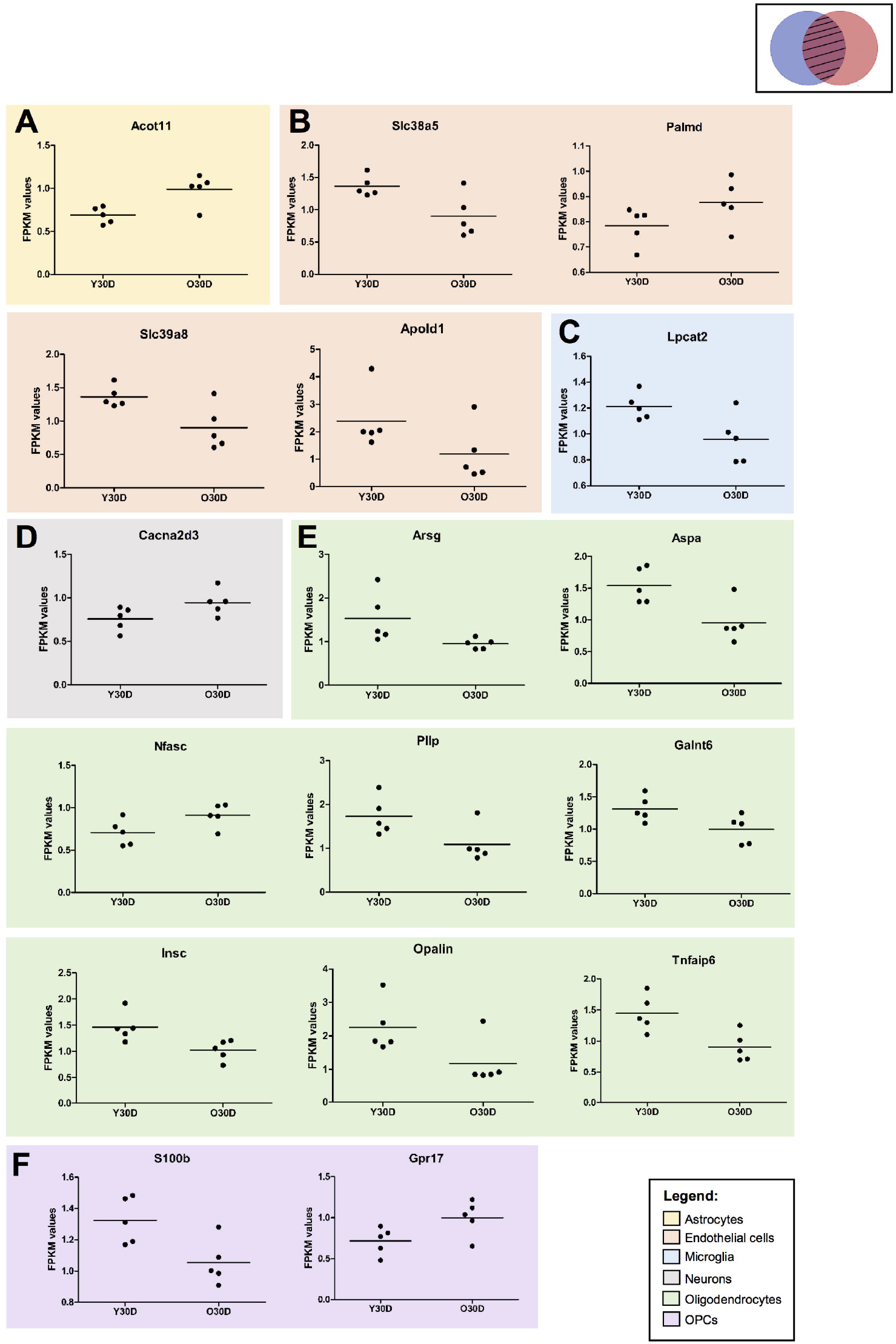
Expression of cell type specific genes overlapping between young 30day BCAS and old Sham animals. (**A**) Astrocyte-associated gene **(B)** endothelial-associated genes **(C)** microglia-associated genes, **(D)** neuron-associated genes, **(E)** oligodendrocyte-associated genes, **(F)** OPC-associated genes. All gene expression values were normalised using the average of the respective controls (i.e. YSham for Y30D and OSham for O30D, respectively). Each dot corresponds to the biological replicates (n=5 for each group). The values are presented as FPKM values obtained through the RNA sequencing results. The solid horizontal lines represent the mean gene expression among the biological replicates.

## Discussion

Chronic cerebral hypoperfusion causes progressive neurodegenerative disease that leads to cognitive decline in the elderly, and is increasingly being recognized as a major health issue. The hippocampus is central for human cognition, memory, executive function, intelligence, path integration and spatial processing (19), all functions that decline with aging and reduced blood supply (38, 39). Understanding how ageing and the associated cerebral hypoperfusion may influence the hippocampal transcriptome may therefore identify mechanisms of impaired function and cognition. The present study is the first to investigate the impact of cerebral hypoperfusion and ageing on the hippocampal transcriptome. The large datasets generated reveal progressive hippocampal transcriptomic responses during hypoperfusion, and somewhat similar changes with ageing. Previous studies have demonstrated that cerebral hypoperfusion leads to white matter hyperintensities and brain atrophy via mechanisms such as energy imbalance, mitochondrial dysfunction, oxidative stress, neuroinflammation and neurovascular dysfunction (40). This study has identified novel genes and signaling pathways in the hippocampus not previously implicated in this condition.

To assess the effect of ageing, we first examined transcriptomic differences between young (3-month old) and old (22-month-old) sham-operated mice. Basal CBF was relatively low in old versus young mice and transcriptomic data showed significant age-related differences in gene expression in the hippocampus. Numerous mitochondrial genes such as *mt-Atp6, mt-Co3, mt-Co2, mt-nd4l* were significantly upregulated in old mice, potentially indicating impaired mitochondrial function. It was similarly reported that expression of other mitochondrial-encoded genes in complexes I, III, IV, and V of the respiratory chain was increased in the mouse brain during ageing (41), and a recent study reported mt-Co2 expression to be increased in neuronal cells stimulated by α-Synuclein (42). The α-Synuclein-induced decline in ATP levels, mitochondrial membrane potential and enhanced reactive oxygen species were reversed by inhibiting mt-Co2 gene expression, thus indicating that increased *mt-Co2* plays a detrimental role (42). The most significantly upregulated DEG, as reflected by the distinct green dot with a high q-value in the volcano plot (**Fig.2A**) was noted to be aldehyde dehydrogenase 1 family member A1 (*Aldh1a1*) as per the heatmap (**Fig.2D**). Expression of *Aldh1a1*, whose role is implicated in neuronal patterns, differentiation and survival (43) is reportedly reduced in Parkinson’s disease (44), perhaps also consistent with a possible disease-causing influence in old age also.

We recently demonstrated that 30 days of cerebral hypoperfusion due to BCAS results in hippocampal neuronal loss (45). In the present study it is noteworthy that, compared with young sham mice, basal CBF was similarly reduced in old sham mice and in young mice subjected to cerebral hypoperfusion for 7 or 30 days. It is therefore plausible that a similar magnitude of chronic CBF reduction could account for overlaps between age- and disease-associated transcriptomic changes in the hippocampus, such as in differential expression of certain mitochondrial and oxidative genes.

We have previously demonstrated that brain expression of cellular stress proteins such as *HSP70* and *GRP78* are greatly diminished in old mice (46). Our transcriptomic data here similarly shows both ageing and cerebral hypoperfusion result in downregulation of neuroprotective heat shock genes such as *Hsph1, Hspa5 (GRP78), Hsp90aa1* and *Hsp90b1*. All proteins encoded by these genes have been shown to protect neurons in multiple neurological conditions (47-49). However, the specific signaling pathways and transcription factors that mediate downregulation of heat shock and other cellular stress proteins are yet to be identified. During ageing, as expected, genes that promote cell death signals were significantly upregulated. Interestingly, this was also seen in young mice following cerebral hypoperfusion. Thus, upregulation of hippocampal genes involved in mitochondrial dysfunction and cell death, and downregulation of cellular protective genes could suggest a convergence of mechanisms resulting from the reduced cerebral blood supply during ageing and BCAS. Given the current evidence, the young mice subjected to cerebral hypoperfusion show convergence in DEGs with the old sham mice, which is suggestive of possible accelerated aging. While further behavioural and histological experiments are required to validate this statement, another study validates our current finding. Wolf and colleagues had shown that young mice subjected to cerebral hypoperfusion and old sham mice were behaviorally analogous in exhibiting decreased social exploration and decreased velocity in open field that were not significantly different from each other (50).

GO term analyses of common downregulated DEGs in the hippocampus of both old and young mice after chronic cerebral hypoperfusion revealed additional ribosome-related genes that when altered may promote disease pathology, including ribonucloprotein complex biogenesis, small and large ribosomal subunits and ribosome. It is well established that loss of RNA homeostasis is a ubiquitous and central feature of neurodegenerative diseases (51). It is noteworthy that telomerase is a ribonucleoprotein complex that maintains telomere length and known to play important roles in neural stem progenitor cell proliferation, neuronal differentiation, survival and neurogenesis (52). Understanding how cerebral hypoperfusion during ageing affects telomerase expression and function will be an important area of future investigation. Ribosomal biogenesis is initiated in the nucleolus and includes the synthesis of ribosomal RNAs, assembly of ribosomal proteins, transportation to the cytoplasm and association of ribosomal subunits (53). Disruption of ribosome biogenesis and decreased expression of small and large ribosomal subunits may promote cell cycle arrest, senescence or apoptosis (53, 54). The relationship between downregulation of ribosomal genes and chronic hypoperfusion-induced brain pathology also requires further investigation.

The cell-type specific analyses, though preliminary, provide an avenue to identify clusters of proteins that play a role in age and disease-associated DEGs. With bioinformatics as a useful tool, this stratification is made possible and it provides a direction of focus for subsequent analyses. Evidence of microglial activation and endothelial dysfunction followed by BBB disruption have been established previously in VaD (1). Hence, studying the different cell types would provide insights into the potential role they might play in driving the pathogenesis of cerebral hypoperfusion.

In summary, this study has performed hippocampal transcriptomic profiling in both ageing and chronic cerebral hypoperfusion that provided important findings. First, progressive transcriptomic changes are induced in both ageing and following chronic cerebral hypoperfusion. Second, interestingly there is an overlap of DEGs and relevant disease pathways between aged and young mice subjected to chronic cerebral hypoperfusion. While limited to a ‘landscape analysis’, our results provide hypothesis-generating information that can be pursued in detail in future studies of hippocampal dysfunction in ageing and models of age-related neurodegenerative diseases such as VaD. Indeed, elucidation of cell-specific pathological mechanisms common to ageing and cerebral hypoperfusion should provide valuable insights for identifying therapeutic targets.

## Materials and Methods

### Experimental Animals and Bilateral Common Carotid Artery Stenosis (BCAS) Mouse Model

All *in vivo* experimental procedures were approved by the National University of Singapore, Singapore Animal Care and Use Committee and performed according to the guidelines set forth by the National Advisory Committee for Laboratory Animal Research (NACLAR), Singapore. All experiments in the manuscript were performed and reported according to ARRIVE (Animal Research: Reporting In Vivo Experiments) guidelines.

Three-months old (Young animals; weighing 24 to 30 grams) and twenty-two months old (Old Animals; weighing 35 to 40 grams) male C57BL/6 mice were obtained from In Vivos, Singapore. Mice were housed in individual cages under standard laboratory conditions. Efforts were taken to minimise the number of animals used and to minimise their suffering during procedures. All mice had free access to food and water *ad libitum*. The experimental groups consist of 9-10 mice in each group. Animals were anaesthetized with isoflurane and subjected to BCAS. BCAS involved using microcoils specially designed for the mice (microcoil specifications: piano wire diameter 0.08mm, internal diameter 0.18mm, coiling pitch 0.5mm, and total length 2.5mm; Sawane Spring Co Ltd, Japan). BCAS was performed by exposing the left and right common carotid arteries (CCAs) one-by-one, freed from their sheaths, and twining a microcoil by rotating it around each CCA. Sham surgeries were performed as controls where the site of surgery was opened and the CCAs were gently touched using the forceps without insertion of microcoils. The site of surgery was subsequently closed using surgical glue, and the mice were observed and taken care of postsurgery until conscious and recovered to freely access food and water *ad libitum*. All animals were euthanized via inhalation of carbon dioxide gas at their respective endpoints after BCAS for subsequent analysis.

### Measurements of Cerebral Blood Flow by Laser Speckle Contrast Imager

High-resolution Laser Speckle Contrast Imager (PSI system, Perimed Inc.) was used to image cerebral blood perfusion and record cerebral blood flow (CBF) before insertion of the microcoils (baseline), immediately after the insertion of the microcoils and finally at their respective end-points of BCAS. Body temperature of the mice was maintained at 37 ± 0.5 °C. The skull of the mice were shaved and exposed by a midline skin incision. The skull was cleaned gently with sterile phosphate buffered saline (PBS) using a cotton applicator. The image area was kept moist and a nontoxic silicon oil was applied on the skull, improving the imaging. Perfusion images were acquired using the PSI system with a 70 mW built-in laser diode for illumination and a 1388 x 1038 pixels CCD camera installed 10 cm above the skull (speed 19 Hz, and exposure time 6 mSec). Analyses of CBF changes were done on the acquired images using a dedicated PIMSoft program (Perimed Inc.).

### Sample Collection and Processing

At the end of each BCAS time point or sham surgeries, the mice were euthanized by administering a lethal dose of carbon dioxide (CO_2_) for inhalation and the brains were harvested. The hippocampi were immediately separated on ice and were stored in a −80°C freezer for RNA sequencing analysis (n = 5 in each experimental group). A separate group of animals were sacrificed for immunoblot analysis (n = 4-5 in each experimental group).

### Total RNA extraction and validation

Total RNA were extracted from frozen brain tissue samples using a micro-tube tissue homogenizer (Bel-Art, Wayne, NJ, USA) and EZ-10 DNAaway RNA extraction mini-prep kit (Bio Basic, Ontario, Canada) following manufacturer’s instructions. Assessment of the integrity and quality of extracted total RNA was performed using agarose gel electrophoresis and Agilent 2100 Bioanalyser (Agilent, Santa Clara, CA, USA); all RNA samples showed RNA integrity numbers above 7, indicating high quality of the extracted total RNA.

### cDNA library preparation and RNA sequencing

The mRNA was purified from total RNA using poly-T oligo-attached magnetic beads and was first fragmented randomly by addition of fragmentation buffer. Subsequently, the first-strand of cDNA was synthesized using a random hexamer primer and M-MuLV reverse transcriptase (RNase H-) (New England BioLabs, Ipswish, MA, USA). Next, DNA polymerase I and RNase H were used to synthesize the second strand. Double-stranded cDNA was purified using AMPure XP beads (Beckman Courter Life Sciences, Indianapolis, IN, USA). Remaining overhangs of the purified doublestranded cDNA were converted into blunt ends via exonuclease/polymerase activities. After adenylation of 3’-ends of DNA fragments, NEBNext adaptor with hairpin loop structure was ligated to prepare for hybridization. In order to select cDNA fragments of preferentially 150-200bp in length, the library fragments were purified with AMPure XP system. Finally, the library was acquired by polymerase chain reaction (PCR) amplification and purification of PCR products by AMPure XP beads. High-throughput sequencing was conducted using HiSeqTM2500 platform (Illumina, San Diego, CA, USA).

### Transcriptome data mapping and differential expression analysis

The RNA sequencing results from the HiSeq system were generated as fasta and quality files, and these files were mapped to the Ensembl-released mouse genome sequence and annotation. Indexes of the reference genome were built using Bowtie V.2.0.6 and paired-end clean reads were aligned to the reference genome using TopHat V.2.0.9 with mismatch parameter limited to 2. For the quantification of gene expression level, HTSeq V.0.6.1 was used to count the read numbers mapped of each gene. Then Reads Per Kilobase of exon model per Million mapped reads (RPKM) of each gene was calculated based on the length of the gene and reads count mapped to the same gene. Differential expression analysis was performed using the DESeq R Package V.1.10.1 and the resulting P-values were adjusted using the Benjamini and Hochberg’s approach for controlling the FDR. Genes with an adjusted P-value lower than 0.05 found by DESeq were assigned as differentially expressed.

### Heatmap generation and enrichment analyses

To create heat maps of differentially expressed genes (DEGs), the R package heatmap3 were used along with the log2Fold-Change output from EdgeR V.3.2.4. To assess the biological significance of gene expression changes, GO and KEGG pathway enrichment analyses were conducted. GO enrichment analysis, focused on biological processes of differentially expressed genes was implemented by the GOseq R package in which gene length bias was corrected. For KEGG pathway enrichment analysis, we used KEGG Orthology-Based Annotation System (KOBAS) software to test the statistical enrichment. GO terms or KEGG pathways with adjusted P-value less than 0.05 were considered significantly enriched by differentially expressed genes.

### Sorting based on cell-type specificity

McKenzie and colleagues (34) had compared murine cell-type specific transcriptome-wide RNA expression data sets from three different groups (35-37). Out of the three mice databases, the top 1000 genes per cell type were selected and ranked according to three cell type associated measures. These include (i) cell-type enrichment which compares if a gene tends to have a higher expression in one cell type compared to the others, (ii) cell-type expression where it compares whether a gene is expressed in a given cell type irrespective of expression levels in the other cell types and (iii) specificity where it compares whether a gene is expressed in only one cell type. Based on the aim of this project, the 1000 genes per cell type were further filtered to the top 100 genes based on specificity and cell enrichment in its respective order.

### Immunoblot Analysis

Brain tissues were homogenized in lysis buffer and then combined with 2 x Laemelli buffer (Bio-Rad Laboratories, Inc., Hercules, CA, USA). Protein samples were then separated on 7.5 to 12.5% v/v sodium dodecyl sulfate (SDS) gels. The SDS-PAGE gels were transferred onto nitrocellulose membranes to probe for proteins. Next, the nitrocellulose membranes were incubated with the following primary antibodies: Mt-ATP6 (Santa Cruz Biotechnology, sc-81886), COX2 (Santa Cruz Biotechnology, sc-514489), COX3 (Santa Cruz Biotechnology, sc-23986), HSP90 (Santa Cruz Biotechnology, sc-13119), GRP78 (Santa Cruz Biotechnology, sc-13539), Calnexin (Santa Cruz Biotechnology, sc-23954), Pink1 (Santa Cruz Biotechnology, sc-517353), PRK8 (Santa Cruz Biotechnology, sc-32282), Fst (Santa Cruz Biotechnology, sc-365003), ActivinA (Novus, NBP1-30928) and β-actin (Sigma-Aldrich, A5441) overnight at 4°C with agitation. Following primary antibody incubation, membranes were washed three times with 1xTBST before incubating with horseradish peroxidase (HRP)-conjugated secondary antibodies (Goat Anti-Rabbit – Cell Signaling Technology, Danvers, MA, USA; Goat Anti-Mouse – Sigma-Aldrich, St. Louis, MO, USA; Goat AntiRat – GE Healthcare Life Sciences, Little Chalfont, UK) for 1 hr at 24 °C with agitation. Following secondary antibody incubation, membranes were washed three times with 1xTBST, each time for 10min. The substrate for HRP, enhanced chemiluminescence (ECL) (Bio-Rad Laboratories, Inc., Hercules, CA, USA) was applied before the membranes were imaged using the ChemiDocXRS+imaging system (Bio-Rad Laboratories, Inc., Hercules, CA, USA). Quantification of proteins was conducted using Image J software (Version 1.46; National Institute of Health, Bethesda, MD, USA), where protein densitometry was expressed relative to the densitometry of the corresponding ß-actin.

## Author Contributions

Study Conception & Design: T.V.A., S.H.B., D.G.J., D.R.H., M.K.P.L., C.L.H.C., G.R.D., K.L.L., and C.G.S; Investigation-Performed Experiment or Data Collection: T.V.A., S.H.B., D.Y.F., and S.S., Data Analysis: S.H.B., S.S., and T.V.A.; Data Interpretation: S.H.B., S.S., and T.V.A.; Writing-Manuscript Preparation and intellectual input: T.V.A., S.H.B., S.S., D.G.J., D.R.H., M.K.P.L., C.L.H.C., G.R.D., K.L.L., and C.G.S; Supervision & Administration: T.V.A., and C.L.H.C.

## References

1) Venkat, P., Chopp, M., and Chen, J. (2015) Models and mechanisms of vascular dementia. Exp Neurol. 272, 97–108.

2) Knopman, D.S., Rocca, W.A., Cha, R.H., Edland, S.D., and Kokmen, E. (2003) Survival study of vascular dementia in Rochester, Minnesota. Arch Neurol, 60, 85–90.

3) Rizzi, L., Rosset, I., and Roriz-Cruz, M. (2014) Global epidemiology of dementia: Alzheimer’s and vascular types. Biomed Res Int. 2014:908–915.

4) Wolters, F.J., and Ikram, M.A. (2019) Epidemiology of Vascular Dementia. Arterioscler Thromb Vasc Biol. 39, 1542–1549.

5) Gorelick, P.B., Scuteri, A., Black, S.E., Decarli, C., Greenberg, S.M., Iadecola, C., Launer, L.J., Laurent, S., Lopez, O.L., Nyenhuis, D., et al. (2011) Vascular contributions to cognitive impairment and dementia: a statement for healthcare professionals from the american heart association/american stroke association. Stroke. 42, 2672–2713.

6) van der Flier, W.M., Skoog, I., Schneider, J.A., Pantoni, L., Mok, V., Chen, C., and Scheltens, P. (2018) Vascular cognitive impairment. Nat Rev Dis Primers, 4, 18003.

7) Parkes, I., Chintawar, S., and Cader, M.Z. (2018) Neurovascular dysfunction in dementia - human cellular models and molecular mechanisms. Clin Sci (Lond). 132, 399–418.

8) Iadecola, C., Duering, M., Hachinski, V., Joutel, A., Pendlebury, S.T., Schneider, J.A., and Dichgans, M. (2019) Vascular Cognitive Impairment and Dementia: JACC Scientific Expert Panel. J Am Coll Cardiol. 73, 3326–3344.

9) van Veluw, S.J., Hilal, S., Kuijf, H.J., Ikram, M.K., Xin, X., Yeow, T.B., Venketasubramanian, N., Biessels, G.J, Chen, C. (2015) Cortical microinfarcts on 3T MRI: Clinical correlates in memory-clinic patients. Alzheimers Dement. 11, 1500–1509.

10) Hilal, S., Sikking, E., Shaik, M.A., Chan, Q.L., van Veluw, S.J., Vrooman, H., Cheng, C.Y., Sabanayagam, C., Cheung, C.Y., Wong, T.Y., Venketasubramanian, N., Biessels, G.J., Chen, C., Ikram, M.K. (2016) Cortical cerebral microinfarcts on 3T MRI: A novel marker of cerebrovascular disease. Neurology. 87, 1583–1590.

11) Hilal, S., Chai, Y.L., van Veluw, S., Shaik, M.A., Ikram, M.K., Venketasubramanian, N., Richards, A.M., Biessels, G.J., Chen, C. (2017) Association Between Subclinical Cardiac Biomarkers and Clinically Manifest Cardiac Diseases With Cortical Cerebral Microinfarcts. JAMA Neurol. 74, 403–410.

12) Ferro, D.A., Mutsaerts, H.J., Hilal, S., Kuijf, H.J., Petersen, E.T., Petr, J., van Veluw, S.J., Venketasubramanian, N., Yeow, T.B., Biessels, G.J., Chen, C. (2020) Cortical microinfarcts in memory clinic patients are associated with reduced cerebral perfusion. J Cereb Blood Flow Metab. 40, 1869–1878.

13) Washida, K., Hattori, Y., and Ihara, M. (2019) Animal Models of Chronic Cerebral Hypoperfusion: From Mouse to Primate. Int J Mol Sci. 20, 6176.

14) Thong, J.Y., Hilal, S., Wang, Y., Soon, H.W., Dong, Y., Collinson, S.L., Anh, T.T., Ikram, M.K., Wong, T.Y., Venketasubramanian, N., Chen, C., Qiu, A. (2013) Association of silent lacunar infarct with brain atrophy and cognitive impairment. J Neurol Neurosurg Psychiatry. 84, 1219–25.

15) Bink, D.I., Ritz, K., Aronica, E., van der Weerd, L., and Daemen, M.J. (2013) Mouse models to study the effect of cardiovascular risk factors on brain structure and cognition. J Cereb Blood Flow Metab. 33, 1666–1684.

16) Ihara, M., Taguchi, A., Maki, T., Washida, K., and Tomimoto, H. (2014) A mouse model of chronic cerebral hypoperfusion characterizing features of vascular cognitive impairment. Methods Mol Biol. 1135, 95–102.

17) Shibata, M., Yamasaki, N., Miyakawa, T., Kalaria, R.N., Fujita, Y., Ohtani, R., Ihara, M., Takahashi, R., and Tomimoto, H. (2007) Selective impairment of working memory in a mouse model of chronic cerebral hypoperfusion. Stroke. 38, 2826–2832.

18) Hattori, Y., Enmi, J., Iguchi, S., Saito, S., Yamamoto, Y., Nagatsuka, K., Iida, H., and Ihara, M. (2016) Substantial Reduction of Parenchymal Cerebral Blood Flow in Mice with Bilateral Common Carotid Artery Stenosis. Sci. Rep. 6, 32179.

19) O’Shea, A., Cohen, R.A., Porges, E.C., Nissim, N.R., and Woods, A.J. (2016) Cognitive aging and the hippocampus in older adults. Frontiers in Aging Neuroscience, 8, 1–8.

20) Raivich, G., and Behrens, A. (2006) Role of the AP-1 transcription factor c-Jun in developing, adult and injured brain. Prog Neurobiol. 78, 347–363.

21) Gao, C., Du, Q., Li, W., Deng, R., Wang, Q., Xu, A., and Shen, J. (2018) Baicalin Modulates APPL2/Glucocorticoid Receptor Signaling Cascade, Promotes Neurogenesis, and Attenuates Emotional and Olfactory Dysfunctions in Chronic Corticosterone-Induced Depression. Mol Neurobiol. 55, 9334–9348.

22) Tretter, Y.P., Hertel, M., Munz, B., ten Bruggencate, G., Werner, S., and Alzheimer, C. (2000) Induction of activin A is essential for the neuroprotective action of basic fibroblast growth factor in vivo. Nat Med. 6, 812–815.

23) Ramming, T., Hansen, H.G., Nagata, K., Ellgaard, L., and Appenzeller-Herzog, C. (2014) GPx8 peroxidase prevents leakage of H2O2 from the endoplasmic reticulum. Free Radic Biol Med. 70, 106–116.

24) Sánchez-Rodríguez, R., Torres-Mena, J.E., Quintanar-Jurado, V., Chagoya-Hazas, V., Rojas Del Castillo, E., Del Pozo Yauner, L., Villa-Treviño, S., Pérez-Carreón, J.I. (2017) Ptgr1 expression is regulated by NRF2 in rat hepatocarcinogenesis and promotes cell proliferation and resistance to oxidative stress. Free Radic Biol Med. 102, 87–99.

25) Newbury, D.F., Winchester, L., Addis, L., Paracchini, S., Buckingham, L.L., Clark, A., Cohen, W., Cowie, H., Dworzynski, K., Everitt, A., et al. (2009) CMIP and ATP2C2 modulate phonological short-term memory in language impairment. Am J Hum Genet. 85, 264–272.

26) Chali, F., Milior, G., Marty, S., Morin-Brureau, M., Le Duigou, C., Savary, E., Blugeon, C., Jourdren, L., and Miles R. (2019) Lipid markers and related transcripts during excitotoxic neurodegeneration in kainate-treated mice. Eur J Neurosci. 50, 1759–1778.

27) Heidbreder, M., Fröhlich, F., Jöhren, O., Dendorfer, A., Qadri, F., and Dominiak, P. (2003) Hypoxia rapidly activates HIF-3alpha mRNA expression. FASEB J. 17, 1541–1543.

28) Zorniak, M., Clark, P.A., and Kuo, J.S. (2015) Myelin-forming cell-specific cadherin-19 is a marker for minimally infiltrative glioblastoma stem-like cells. J Neurosurg. 122, 69–77.

29) McGahan, L., Hakim, A.M., Nakabeppu, Y., and Robertson, G.S. (1998) Ischemia-induced CA1 neuronal death is preceded by elevated FosB and Jun expression and reduced NGFI-A and JunB levels. Brain Res Mol Brain Res. 56, 146–161.

30) Hughes, P.E., Alexi, T., Walton, M., Williams, C.E., Dragunow, M., Clark, R.G., and Gluckman, P.D. (1999) Activity and injury-dependent expression of inducible transcription factors, growth factors and apoptosis-related genes within the central nervous system. Prog Neurobiol. 57, 421–50.

31) Lueptow, L.M., Devi, L.A., and Fakira, A.K. (2018) Targeting the Recently Deorphanized Receptor GPR83 for the Treatment of Immunological, Neuroendocrine and Neuropsychiatric Disorders. Prog Mol Biol Transl Sci. 159, 1–25.

32) Cheng, Y., Yang, C., Zhao, J., Tse, H.F., and Rong, J. (2015) Proteomic identification of calcium-binding chaperone calreticulin as a potential mediator for the neuroprotective and neuritogenic activities of fruit-derived glycoside amygdalin. J Nutr Biochem. 26, 146–154.

33) Pickrell, A.M., and Youle, R.J. (2015) The roles of PINK1, parkin, and mitochondrial fidelity in Parkinson’s disease. Neuron. 85, 257–273.

34) McKenzie, A.T., Wang, M., Hauberg, M.E., Fullard, J.F., Kozlenkov, A., Keenan, A., Hurd, Y.L., Dracheva, S., Casaccia, P., Roussos, P., and Zhang, B. (2018) Brain Cell Type Specific Gene Expression and Co-expression Network Architectures. Sci Rep. 8, 8868.

35) Tasic, B., Menon, V., Nguyen, T.N., Kim, T.K., Jarsky, T., Yao, Z., Levi, B., Gray, L.T., Sorensen, S.A., Dolbeare, T., et al. (2016) Adult mouse cortical cell taxonomy revealed by single cell transcriptomics. Nat Neurosci. 19, 335–346.

36) Zeisel, A., Muñoz-Manchado, A.B., Codeluppi, S., Lönnerberg, P., La Manno, G., Juréus, A., Marques, S., Munguba, H., He, L., Betsholtz, C., et al. (2015) Brain structure. Cell types in the mouse cortex and hippocampus revealed by singlecell RNA-seq. Science. 347, 1138–1142.

37) Zhang, Y., Sloan, S.A., Clarke, L.E., Caneda, C., Plaza, C.A., Blumenthal, P.D., Vogel, H., Steinberg, G.K., Edwards, M.S., Li, G., et al. (2015) Purification and Characterization of Progenitor and Mature Human Astrocytes Reveals Transcriptional and Functional Differences with Mouse. Neuron. 89, 37–53.

38) Bettio, L.E.B., Rajendran, L., and Gil-Mohapel, J. (2017) The effects of aging in the hippocampus and cognitive decline. Neurosci Biobehav Rev. 79, 66–86.

39) Park, J.H., Hong, J.H., Lee, S.W., Ji, H.D., Jung, J.A., Yoon, K.W., Lee, J.I., Won, K.S., Song, B.I., and Kim, H.W. (2019) The effect of chronic cerebral hypoperfusion on the pathology of Alzheimer’s disease: A positron emission tomography study in rats. Sci Rep. 9, 14102.

40) Aliev, G., Obrenovich, M.E., Smith, M.A., and Perry, G. (2003) Hypoperfusion, mitochondria failure, oxidative stress, and Alzheimer disease. J Biomed Biotechnol. 3, 162–163.

41) Manczak, M., Jung, Y., Park, B.S., Partovi, D., and Reddy, P.H. (2005) Time course of mitochondrial gene expressions in mice brains: implications for mitochondrial dysfunction, oxidative damage, and cytochrome c in aging. J Neurochem. 92, 494–504.

42) Danyu, L., Yanran, L., Xiuna, J., Ying, C., Sudan, P., Tianen, Z., Zhifen, Z., Dezhi, Z., Kaixun, H., Yingyu, X., and Enxiang T. (2019) α-Synuclein induced mitochondrial dysfunction via cytochrome c oxidase subunit 2 in SH-SY5Y cells. Exp Cell Res. 378, 57–65.

43) Sgobio, C., Wu, J., Zheng, W., Chen, X., Pan, J., Salinas, A.G., Davis, M.I., Lovinger, D.M., and Cai, H. (2017) Aldehyde dehydrogenase 1-positive nigrostriatal dopaminergic fibers exhibit distinct projection pattern and dopamine release dynamics at mouse dorsal striatum. Sci Rep. 7, 5283.

44) Wey, M.C., Fernandez, E., Martinez, P.A., Sullivan, P., Goldstein, D.S., and Strong, R. (2012) Neurodegeneration and motor dysfunction in mice lacking cytosolic and mitochondrial aldehyde dehydrogenases: implications for Parkinson’s disease. PloS one, 7, e31522.

45) Poh, L., Fann, D.Y., Wong, P., Lim, H.M., Foo, S.L., Kang, S.W., Rajeev, V., Selvaraji, S., Vinaya, R.I., et al. (2020) AIM2 Inflammasome Mediates Hallmark Neuropathological Alterations and Cognitive Impairment in a Mouse Model of Vascular Dementia. bioRxiv 135228

46) Arumugam, T.V., Phillips, T.M., Cheng, A., Morrell, C.H., Mattson, M.P., and Wan, R. (2010) Age and energy intake interact to modify cell stress pathways and stroke outcome. Ann Neurol. 67, 41–52.

47) Scior, A., Buntru, A., Arnsburg, K., Ast, A., Iburg, M., Juenemann, K., Pigazzini, M.L., Mlody, B., Puchkov, D., Priller, J, et al. (2018) Complete suppression of Htt fibrilization and disaggregation of Htt fibrils by a trimeric chaperone complex. EMBO J. 37, 282–299.

48) O’Brien, R., DeGiacomo, F., Holcomb, J., Bonner, A., Ring, K.L., Zhang, N., Zafar, K., Weiss, A., Lager, B., Schilling, B., et al. (2015) Integration-independent Transgenic Huntington Disease Fragment Mouse Models Reveal Distinct Phenotypes and Life Span in Vivo. J Biol Chem. 290, 19287–19306.

49) Zhang, M., Qian, C., Zheng, Z.G., Qian, F., Wang, Y., Thu, P.M., Zhang, X., Zhou, Y., Tu, L., Liu, Q., et al. (2018) Jujuboside A promotes Aβ clearance and ameliorates cognitive deficiency in Alzheimer’s disease through activating Axl/HSP90/PPARγ pathway. Theranostics. 8, 4262–4278.

50) Wolf, G., Lotan, A., Lifschytz, T., Ben-Ari, H., Kreisel Merzel, T., Tatarskyy, P.,… & Lerer, B. (2017). Differentially Severe Cognitive Effects of Compromised Cerebral Blood Flow in Aged Mice: Association with Myelin Degradation and Microglia Activation. Frontiers in aging neuroscience, 9(191), 1–13.

51) Pellizzoni L. (2007) Chaperoning ribonucleoprotein biogenesis in health and disease. EMBO reports. 8, 340–345.

52) Liu, M.Y., Nemes, A., and Zhou, Q.G. (2018) The Emerging Roles for Telomerase in the Central Nervous System. Front Mol Neurosci. 11, 160.

53) Turi, Z., Lacey, M., Mistrik, M., and Moudry, P. (2019) Impaired ribosome biogenesis: mechanisms and relevance to cancer and aging. Aging (Albany NY). 11, 2512–2540.

54) Narla, A., and Ebert, B.L. (2010) Ribosomopathies: human disorders of ribosome dysfunction. Blood. 115, 3196–3205.

