## Supplementary Figure Legends for "Hippocampal Transcriptome Profiling Reveals Common Disease Pathways in Chronic Hypoperfusion and Ageing"

**Supplementary figure legend**

**Supplementary Figure 1**| Hierarchical clustering analysis of all 11 differentially expressed genes in O30D compared to OSham mice. There were no significant DEGs in O7D compared to OSham mice and hence is not represented in the heatmap. Upregulated gene expression is represented in red and downregulated gene expression in blue. The colour scale represents the log10 (average FPKM+1) value.

**Supplementary Figure 2** | Significant GO terms in Y7D vs YSham mice (q < 0.05). The horizontal axis represents the -log10 scale for q-value of each GO term.

**Supplementary Figure 3** | KEGG pathway enrichment analysis between Y7D and YSham mice. The size of the dots represents the gene ratio involvement and the colour represents the significance. Red represents most significant and purple represents the least significant with the gradient showing the q-value range.

**Supplementary Figure 4** | Significant GO terms in Y30D vs YSham mice (q < 0.05). The horizontal axis represents the -log10 scale for q-value of each GO term.
