## Supplementary figures and images for "Hippocampal Transcriptome Profiling Reveals Common Disease Pathways in Chronic Hypoperfusion and Ageing"

### S.Figure 1

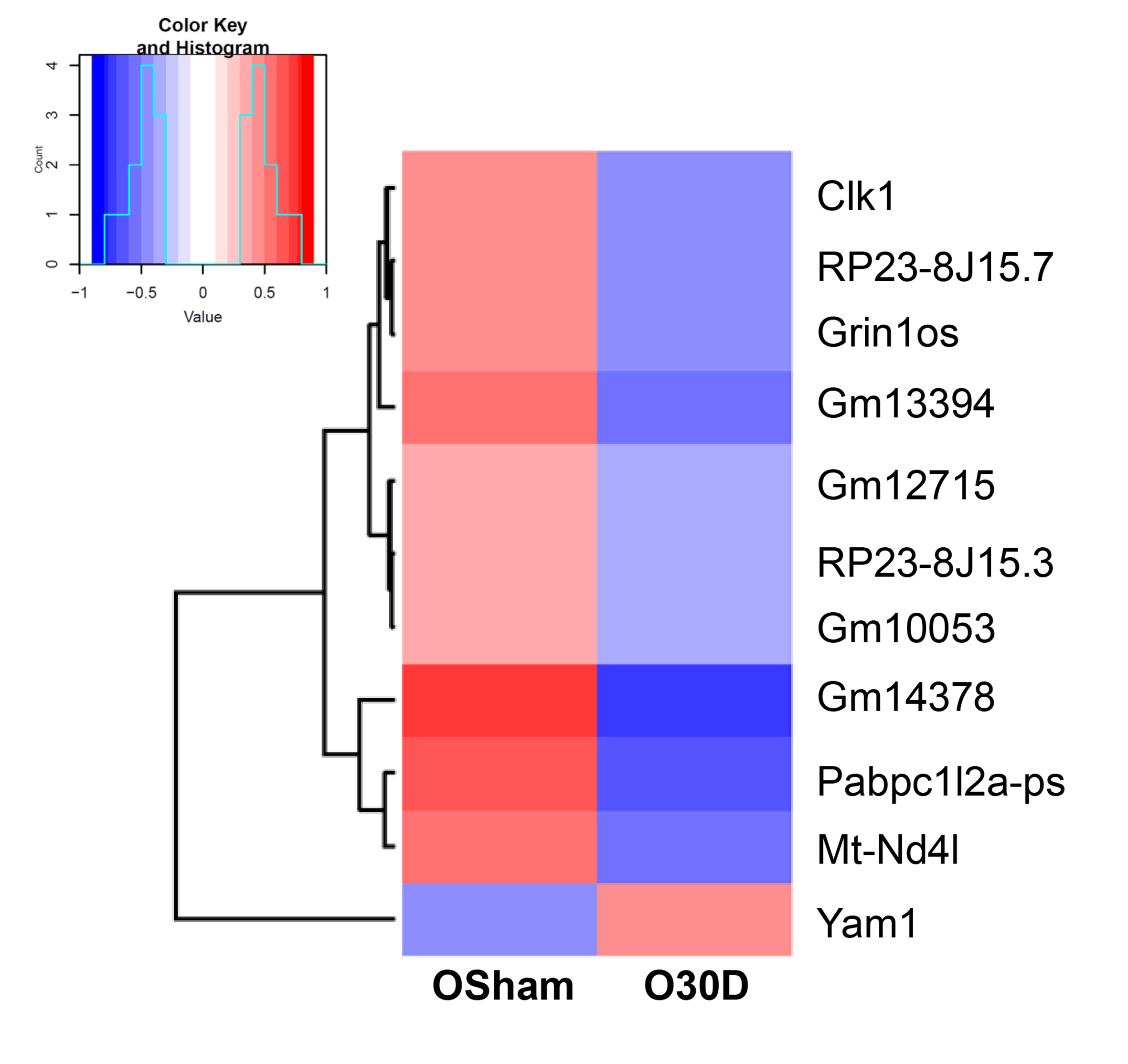

### S.Figure 2

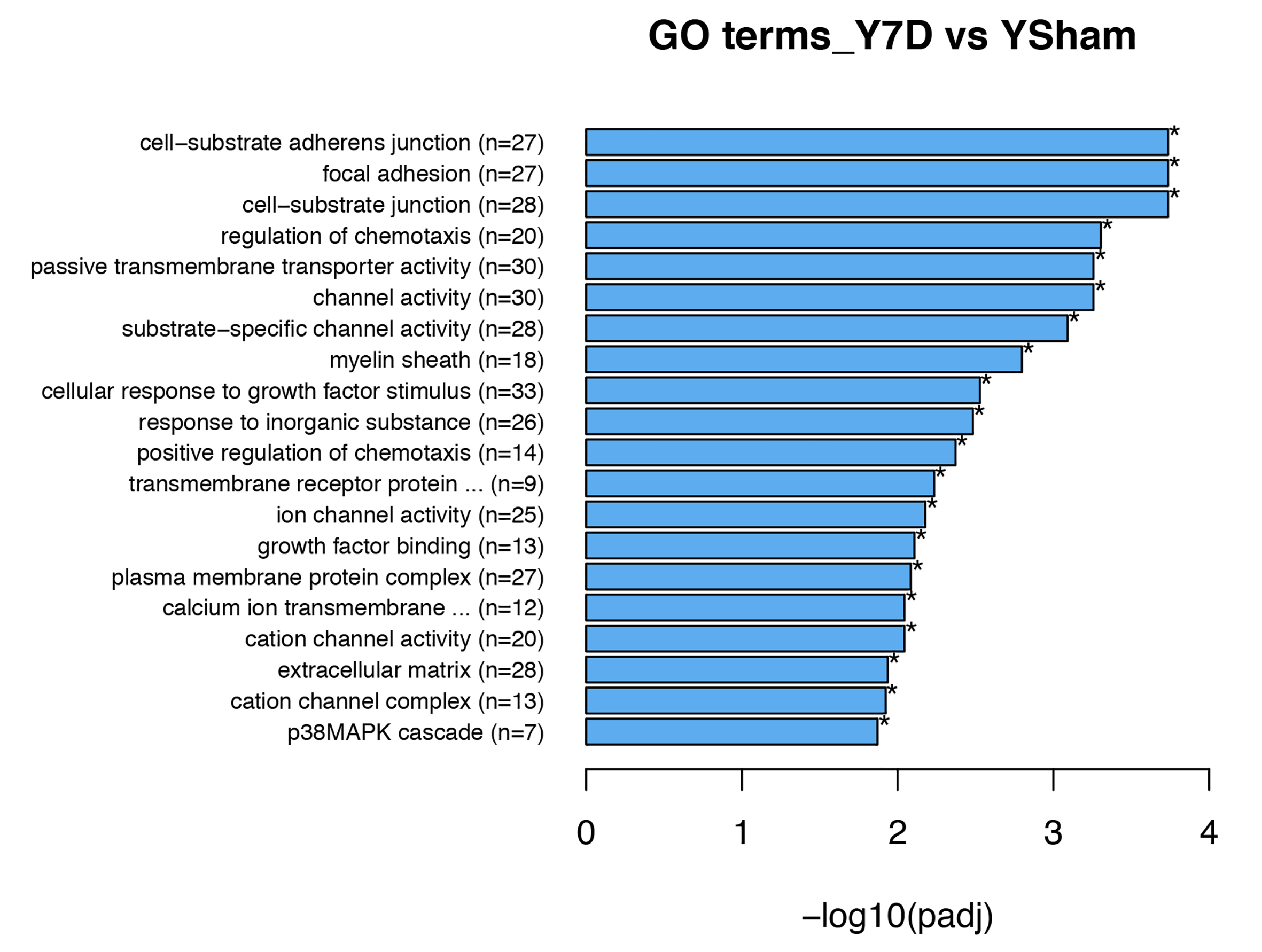

### S.Figure 3

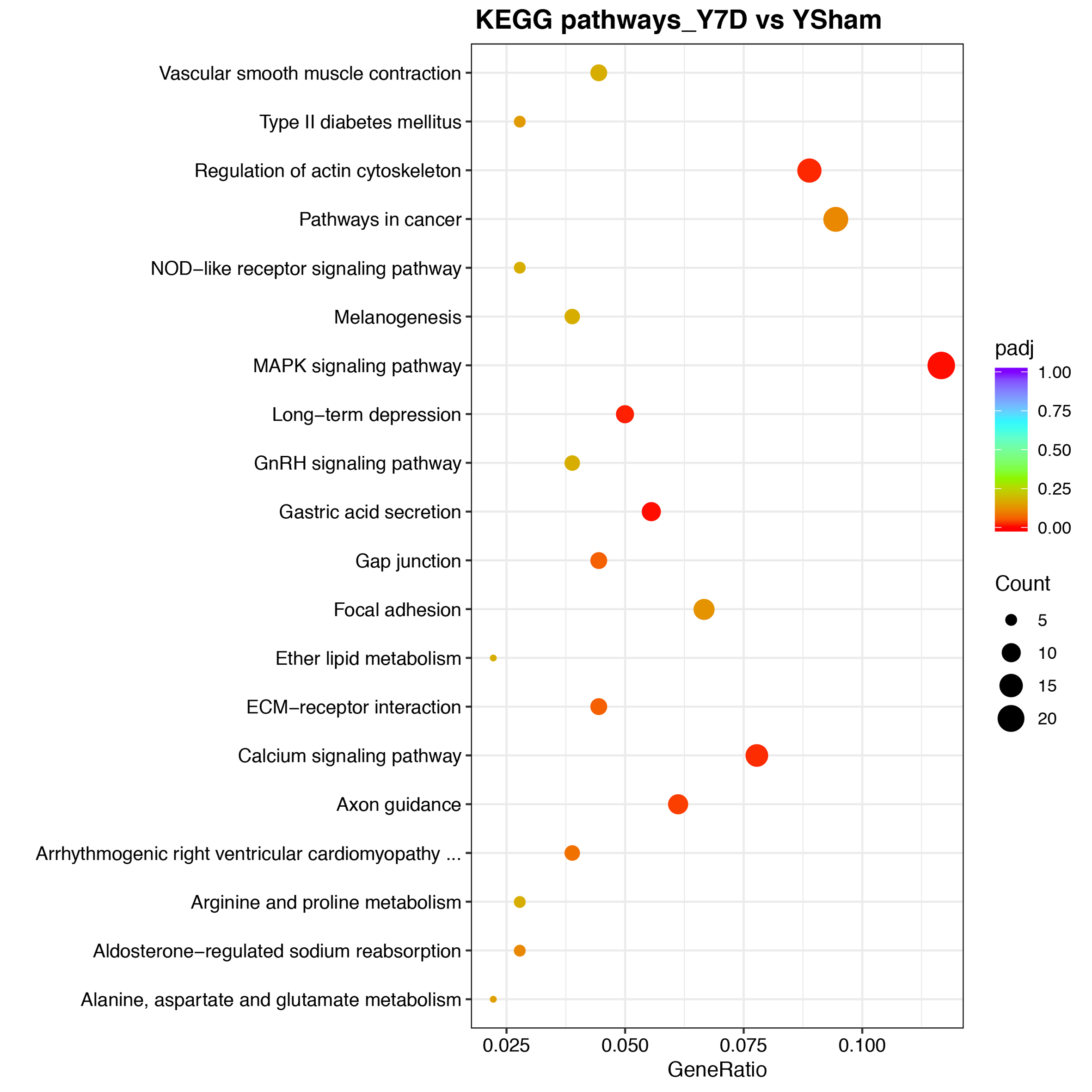

### S.Figure 4

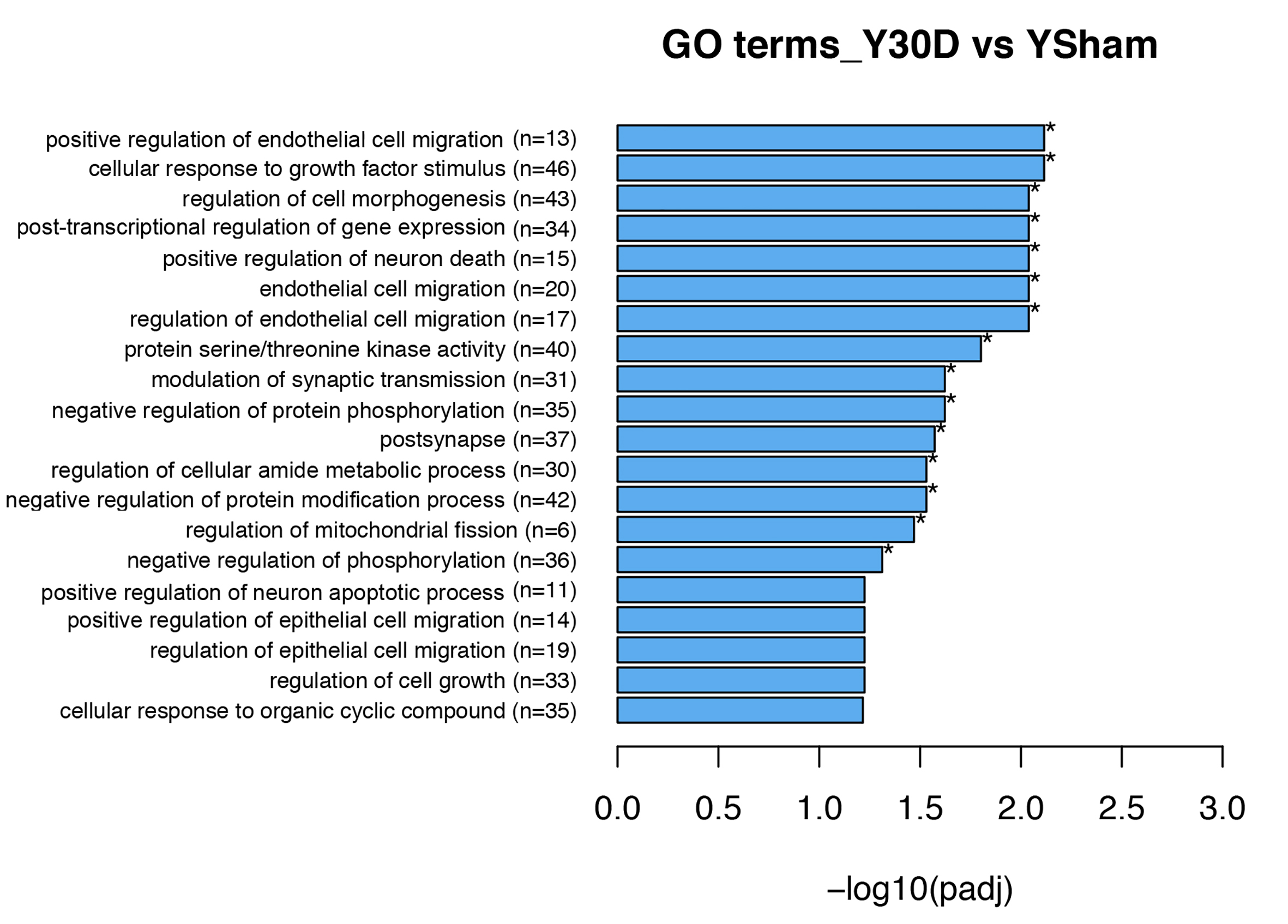
